# eCovSens-Ultrasensitive Novel In-House Built Printed Circuit Board Based Electrochemical Device for Rapid Detection of nCovid-19 antigen, a spike protein domain 1 of SARS-CoV-2

**DOI:** 10.1101/2020.04.24.059204

**Authors:** Subhasis Mahari, Akanksha Roberts, Deepshikha Shahdeo, Sonu Gandhi

**Author notes:** Corresponding author- Dr Sonu Gandhi. Authors contributed equally (Subhasis Mahari, Akanksha Roberts).

## Abstract

Severe acute respiratory syndrome coronavirus 2 (SARS-CoV-2 or nCovid-19) outbreak has become a huge public health issue due to its rapid transmission and global pandemic. Currently, there are no vaccines or drugs available for nCovid-19, hence early detection is crucial to help and manage the outbreak. Here, we report an in-house built biosensor device (eCovSens) and compare it with a commercial potentiostat for the detection of nCovid-19 spike antigen (nCovid-19Ag) in spiked saliva samples. A potentiostat based sensor was fabricated using fluorine doped tin oxide electrode (FTO) with gold nanoparticle (AuNPs) and immobilized with nCovid-19 monoclonal antibody (nCovid-19Ab) to measure change in the electrical conductivity. Similarly, eCovSens was used to measure change in electrical conductivity by immobilizing nCovid-19 Ab on screen printed carbon electrode (SPCE). The performances of both sensors were recorded upon interaction of nCovid-19Ab with its specific nCovid-19Ag. Under optimum conditions, the FTO based immunosensor and eCovSens displayed high sensitivity for detection of nCovid-19Ag, ranging from 1 fM to 1 μM. Our in-house developed device can successfully detect nCovid-19Ag at 10 fM concentration in standard buffer that is in close agreement with FTO/AuNPs sensor. The limit of detection (LOD) was found to be 90 fM with eCovSens and 120 fM with potentiostst in case of spiked saliva samples. The proposed portable eCovSens device can be used as a diagnostic tool for the rapid (within 10-30 s) detection of nCovid-19Ag traces directly in patient saliva in a non-invasive manner.

## 1. Introduction

In December 2019, an outbreak of pneumonia was observed in Wuhan, China and the causative pathogen was identified as a new type of coronavirus (Chan et al., 2020; Chen et al., 2020; Ren et al., 2020; Wu et al., 2020; Zhu et al., 2020) and named as 2019 novel coronavirus (2019‐ nCoV) by the World Health Organization (WHO). By comparing taxonomy, phylogeny, and established practice, the Coronavirus Study Group of the International Committee on Taxonomy of Viruses established this virus to be related to severe acute respiratory syndrome coronavirus (SARS‐CoV) and renamed it as SARS‐CoV‐2 (Gorbalenya et al., 2020). SARS-CoV-2 is a single positive strand RNA virus consisting of four structural proteins including spike (S), envelope (E), matrix (M), and nucleocapsid (N) proteins and is responsible for respiratory tract illness in humans (Li et al., 2020a). According to recent studies, SARS-CoV-2 utilizes angiotensin converting enzyme 2 (ACE2) as a receptor for cellular entry making the use of spike (S) protein with high affinity towards ACE2 (Tian et al., 2020; Wrapp et al., 2020).

As of 24^th^ April, 2020, the total number of nCovid-19 cases in the world have surpassed 2 million with 27,25,920 fatalities globally. With more than half the world under lock-down in order to contain the spread of this pandemic, with no immediate vaccine available, it has become the need of the hour to develop rapid detection methods to diagnose nCovid-19 in both symptomatic as well as asymptomatic patients (Mizumoto et al., 2020) that would enable early mitigation. For early diagnosis, chest computed tomography (CT) (Bai et al., 2020; Bernheim et al., 2020; Li and Xia, 2020; Pan et al., 2020a) was used whereas in the analytical stage, real-time reverse-transcriptase polymerase chain reaction (RT-PCR) (Chu et al., 2020; Corman et al., 2020; Lan et al., 2020; Loeffelholz and Tang, 2020) remains the standard test for the etiologic diagnosis of SARS-CoV-2 (Ai et al., 2020; Fang et al., 2020). Recently, antibody and CRISPR based techniques are being introduced as supplemental tools for rapid diagnosis (Li et al., 2020b; Ding et al., 2020; Yang et al., 2020).

Biosensors have the advantage of being sensitive, specific, stable, easy to use, require less sample size, time, portable, and most importantly can be customised to detect the target analyte of interest. Immunosensors can be used to detect toxins (Kasoju et al., 2020a), narcotic drugs (Gandhi et al., 2018, Mishra et al., 2018; Singh et al., 2017; Tey et al., 2010) viruses (Kerry et al., 2019) by use of different bioreceptors such as deoxyribonucleic acid (DNA) (Jiang et al., 2005; Labuda et al., 2009), enzymes (Ilangovan et al., 2006; Jawaheer et al., 2003), peptides (Gandhi et al., 2016), aptamers (Kasoju et al., 2020b), antibody (Islam et al., 2019; Roberts et al., 2019). Electrochemical biosensors are considered as a reliable tool for infectious disease detection as they remain unaffected by sample absorbance or turbidity (Bakker, 2004). In order to increase the sensitivity of electrochemical biosensors, nanomaterials are often made use of a signal amplifiers such as graphene (Islam et al., 2019), and AuNPs (Pingarrón et al., 2008).

In this work, we have developed an in-house built device named as eCovSens using an SPCE electrode and compared it with a potentiostat using FTO electrode. The comparison was made in terms of sensitivity, specificity, time of detection, sample volume, portability, and stability. Here, FTO electrodes have been preferred over indium tin oxide (ITO) electrodes due to its high electrical conductivity, chemical stability under atmospheric conditions, high tolerance towards physical abrasions and cost effectiveness (Roberts et al., 2019). AuNPs were selected as the signal amplifiers due to its high conductivity, biocompatibility, stability, and size related electronic properties (Talan et al., 2018). AuNPs were drop casted onto the FTO electrode and nCovid-19 Ab was immobilized to detect the presence of nCovid-19 spike Ag. All immobilization steps were characterised using physicochemical methods such as UV-Vis Spectroscopy, Transmission electron microscopy (TEM), Dynamic light scattering spectroscopy (DLS), Fourier transform infra-red spectroscopy (FT-IR), Cyclic Voltammetry (CV), Differential Pulse Voltammetry (DPV). In case of eCovSens, nCovid-19 Ab was immobilized onto SPCE for detection. The LOD of nCovid-19 Ag was determined at 10 fM (in-house built device) which was in close proximity with potentiostat with the range of concentrations from 1 fM to 1 μM in standard buffer. We concluded that the eCovSens is ultrasensitive, highly specific, provides rapid results within 10-30 s, requires only 20 μL sample volume, and can be carried to the bedside and is stable upto one month if compared with potentiostat that requires a laboratory set up and 100 μl sample volume. Furthermore, eCovSens device can be customised to any target analyte, and can also have other future applications for detection of various other ailments.

## 2. Materials and Methods

### 2.1. Reagents and Apparatus

Sodium dihydrogen phosphate-1-hydrate (Na_2_HPO_4_.H_2_O) was acquired from Merck (Mumbai, India). Potassium dihydrogen orthophosphate (KH_2_PO_4_), sodium carbonate anhydrous (Na_2_CO_3_), sodium bicarbonate (NaHCO_3_), sodium citrate tribasic dehydrate (C_6_H_5_Na_3_O_7_.2H_2_O), potassium ferrocynaide (K_4_Fe(CN)_6_·3H_2_O) and potassium ferricyanide (C_6_N_6_FeK_3_) were procured from Sisco Research Laboratories (SRL, India). Potassium chloride (KCl) and sodium chloride (NaCl) were obtained from CDH (New Delhi, India). Carbon coated copper TEM grids were acquired from Ted Pella Inc. (Redding, Canada). The FTO electrodes and gold(III) chloride (Au_2_Cl_6_) were purchased from Sigma-Aldrich (India) while the SPCE from Zensor (Texas, USA). nCovid-19 Ag (Spike S1 protein) and nCovid-19 Ab were procured from ProSci (California, USA). Japanese Encephalitis Virus (JEV), Human Immunodeficiency Virus (HIV) and Avian Influenza Virus (AIV) Ag were obtained from The Native Antigen Company (Oxford, UK). Aurdino software has been used in the in-house built device and the hardware include a printed Circuit Board (PCB), the encoder and decoder, adapter, bluetooth module, Op amplifier, resistors, and transistors. All chemicals, solvents and reagents used were of high quality analytical grade unless stated otherwise and all solutions were prepared in double distilled water.

### 2.2. Instrumentation

UV-Vis and FT-IR spectra were acquired on Systonic S-924 Single-Beam UV-Vis Spectrophotometer (Delhi, India) and Thermo Scientific-Nicolet iS50 FT-IR (Bangalore, India) respectively. Changes in hydrodynamic diameter and zeta potential of each immobilization step were observed using Anton-Paar Litesizer 500 Particle Analyzer DLS (Gurgaon, India). Morphology and size were observed in TEM images that were obtained by JEOL-JEM 2010 operated at an accelerating voltage of 200 kV. The CV and DPV measurements were performed with Metrohm Autolab-PGSTAT-101 (Chennai, India) driven by Nova 2.1 software. Connecting of wires in the in-house built device were done using a soldering device and a Mastech Group multimeter was used to check the proper flow of current in the circuit during fabrication. All experiments were performed at room temperature (RT) (25 °C) unless stated otherwise.

### 2.3. Synthesis of AuNPs and its labelling with nCovid-19 Ab

AuNPs were synthesized using Turkevich (Turkevich et al., 1951) and Frens (Frens, 1973) heat-reflux citrate reduction method. For the synthesis of AuNPs, gold chloride (0.01 mL, 10%) was added to milli-Q water and heated until the mixture began to boil. 1 mL of 1 % sodium citrate tribasic was immediately added to the boiling solution, that resulted in gradual change in the colour from yellow to dark blue and finally to wine red. The colloidal solution cooled and stored at 4 °C until further use. For labelling of nCovid-19 Ab with AuNPs, 90 μg nCovid-19 Ab was added drop wise to 1 mL of AuNPs solution in Phosphate Buffer (PB) (20 mM, pH 7.5). The AuNPs/nCovid-19 Ab mix was allowed to react overnight (O/N) at 4 °C and centrifuged at 12,000 rpm for 30 min at 4 °C to remove any excess unbound Ab. The AuNPs/nCovid-19 Ab conjugate was resuspended in PB (20 mM, pH 7.5) for further studies.

### 2.4. Characterisation of AuNPs and AuNPs/nCovid-19 Ab

Various physicochemical methods were used to confirm the labelling of nCovid-19 Ab with AuNPs. UV-Vis spectra were observed in the range of 200-800 nm with a step of 0.1 nm and scanning speed of 20 nm/s for AuNPs and AuNPs/nCovid-19 Ab conjugate. The hydrodynamic diameter and zeta potential of both the AuNPs and AuNPs/nCovid-19 Ab conjugate were obtained from the DLS at 200 kHz for photon counting, scattering angle 90 °C, and temperature 24 ± 2 °C. The hydrodynamic diameter calculated based on Stokes–Einstein equation as water was considered as the continuous phase (water viscosity = 0.911–0.852 mPa/s, diffusion coefficient of AuNPs = 6.89 × 10^−9^ to 5.30 × 10^−8^ cm^2^/s) (Talan et al., 2018). FT-IR spectra were taken in the range of 1000-4000 cm^−1^ to determine the changes in bonds/functional groups. Morphology and size were determined by drop casting the AuNPs and AuNPs/nCovid-19 Ab samples on carbon coated copper TEM grids.

### 2.5. Fabrication of FTO electrode with AuNPs/nCovid-19 Ab

The FTO (3 cm × 5 cm) electrode was made up of glass coated with fluorine doped tin oxide. 200 μl of AuNPs were drop casted on the surface of FTO electrode and completely dried for 48 h at 4 °C. 40 μl of nCovid-19 Ab (1 μg/mL) was immobilised on different FTO/AuNPs electrode further for 24 h at 4 °C.

### 2.6. Characterisation, optimisation, and testing of FTO/AuNPs/nCovid-19 Ab with nCovid-19 Ag

Electrochemical characterisation of the FTO/AuNPs/nCovid-19 Ab was done by using FTO as a working electrode, and Ag/AgCl as a reference electrode. This was done by sweeping the potential from −0.001 kV to 0.001 kV in K_3_[Fe(CN)_6_]/K_4_[Fe(CN)_6_] (1:1) solution containing 100 mM KCl. In order to obtain maximum sensing signal, various factors such as Ab concentration, temperature, pH, and response time, were optimised by comparing CV/DPV. The nCovid-19 Ag concentration was prepared in the range of 1 fM to 1 μM (in 1X PBS, pH 7.4) and LOD was determined. The application of the fabricated FTO/AuNPs/nCovid-19 Ab sensor was also evaluated for its degree of sensitivity in saliva samples spiked with nCovid-19 Ag (120 fM). Furthermore, the stability and repeatability of the fabricated FTO/AuNPs/ nCovid-19 Ab was also evaluated over a period of 4 weeks. The specificity was determined by analyzing cross-reactivity against other viral Ag including HIV, JEV, and AIV.

### 2.7. Fabrication of the novel in-house built electrochemical device

The designed in-house electrochemical device can be called an embedded system that is a combination of elements, software package, and other mechanical/distinct components designed to perform a selected operation. Each embedded system consists of a custom-made hardware designed around a central processing unit (CPU) and this hardware additionally contains memory chips onto which the software package is loaded. The software package residing on the microchip is additionally known as the computer code. The embedded system design is often diagrammatical as a stratified design. The building blocks of the embedded system include a central processing unit (CPU), pair of memory storages (read-only memory (ROM), and random access memory (RAM)), input devices, output devices, communication interfaces, and application-specific electronic equipment.

**Flowchart 1.**
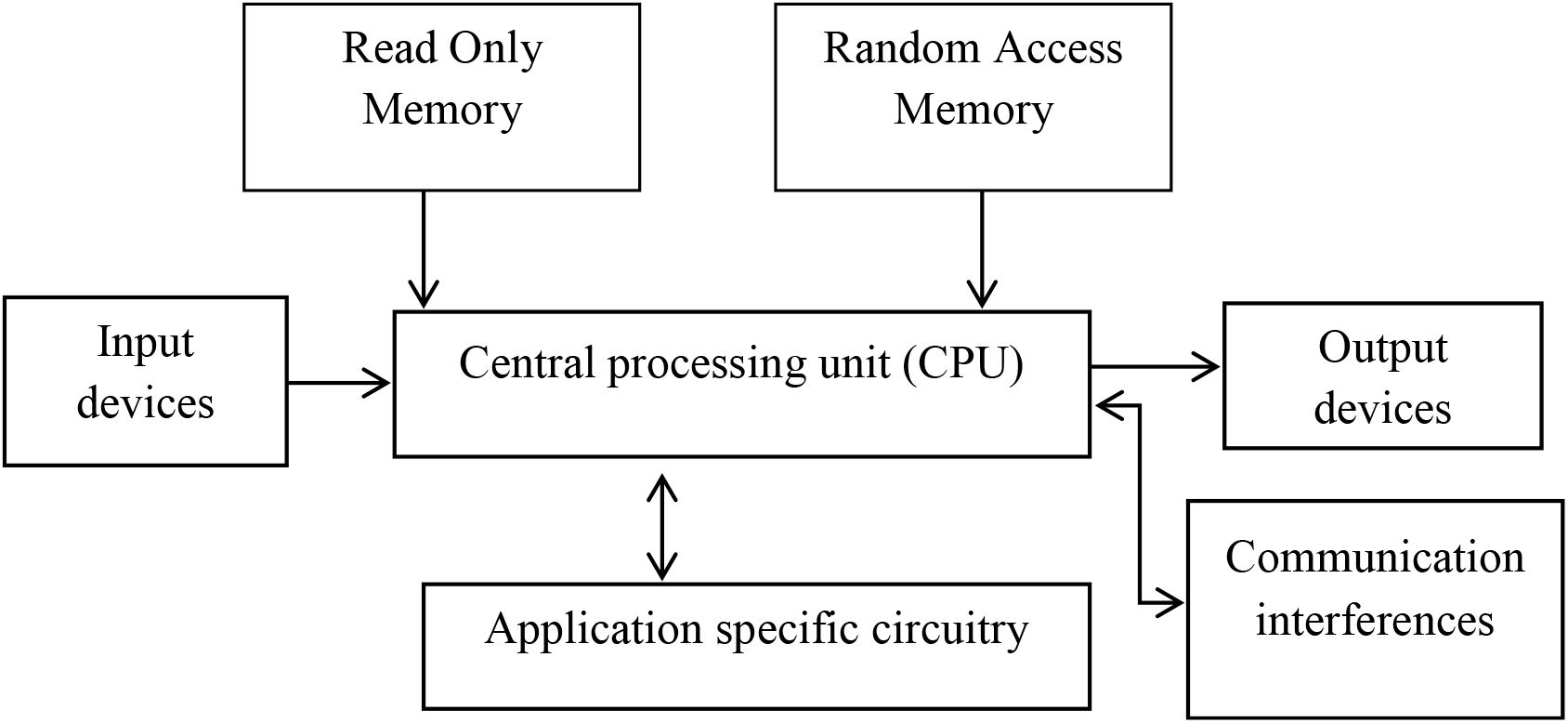
The hardware architecture of the embedded system with inputs and output devices and communication interferences

Flowchart 1 showed the different components of the hardware design of the in-house device embedded system where the CPU acts as the associate in nursing interface between input and output devices. Additionally, it consists of RAM, and storage recollections that are able to store temporary, and permanent information. Communication system acts as an associate in the nursing interface between CPU, and different parts of the embedded system. Fabrication of the device was carried out as follows: Using the free version of Aurdino integrated development environment (IDE), we designed the biosensor in two steps: (i) the circuit diagram using PROTEUS design software as the stimulation tool, and (ii) the route of the wires, and layout of the components on a two-layer printed circuit board (PCB). The components were surface-mounted onto the PCB. The firmware of the microcontroller (RFduino) was written, using EMBEDDED C programming. To load the firmware onto the microcontroller, a universal serial bus (USB) was attached to the contact pads on the PCB, and connected the other end to the terminals of a USB shield for Aurdino. The socket was created and connected with SPCE, and a rechargeable to the contact pads.

### 2.8. Specificity and cross-reactivity studies of the fabricated in-house built device

Optimisation of all the parameters such as Ab concentration, buffer conditions were done before carrying out the detection (data not shown). The SPCE electrode was inserted into the device and change in voltage was measured in mV. For this, 0.02 μg of nCovid-19 Ab (20 μL) was immobilized on the electrode via passive adsorption. Different concentrations of nCovid-19 Ag were applied onto the electrode ranging from 1 fM to 1 μM to analyze the sensitivity of the device. Cross-reactivity study was done using AIV Ag at similar concentrations as of nCovid-19 Ag. The nCovid-19 Ag (90 fM) was spiked into saliva and monitored for its sensitivity. The electrode was washed with PBS after each step, and values were noted immediately (in 10 to 30 s) after addition of buffer/antibody/antigen. Furthermore, the stability and repeatability of the designed sensor was evaluated on the same SPCE/nCovid-19 Ab electrode over a period of 4 weeks at 7 day intervals.

## 3. Results and Discussion

### 3.1. Design and principle of the fabricated FTO/AuNPs/nCovid-19 Ab sensor

Scheme 1 elucidates the mechanism of sensing and fabrication of the developed FTO immunosensor integrated with AuNPs and nCovid-19 Ab. AuNPs act as a catalyst and amplify the electrochemical signal by enhancement of electrical conductivity. Presence of AuNPs serves as a platform for the attachment of nCovid-19 Ab by electrostatic interactions or simple physisorption. Addition of nCovid-19 Ag on the FTO/AuNPs/nCovid-19 Ab modified electrode led to change in electrical current. The major phenomenon that lies behind, is the orientation and polarity of the protein molecules that plays a crucial role in electron transfer from the electrode surface. The developed immunosensor effectively combined the beneficial features of FTO electrode, AuNPs, and the highly specific immunological interaction between nCovid-19 Ab and its nCovid-19 Ag, that enabled quick and effective response.

**Scheme 1.**
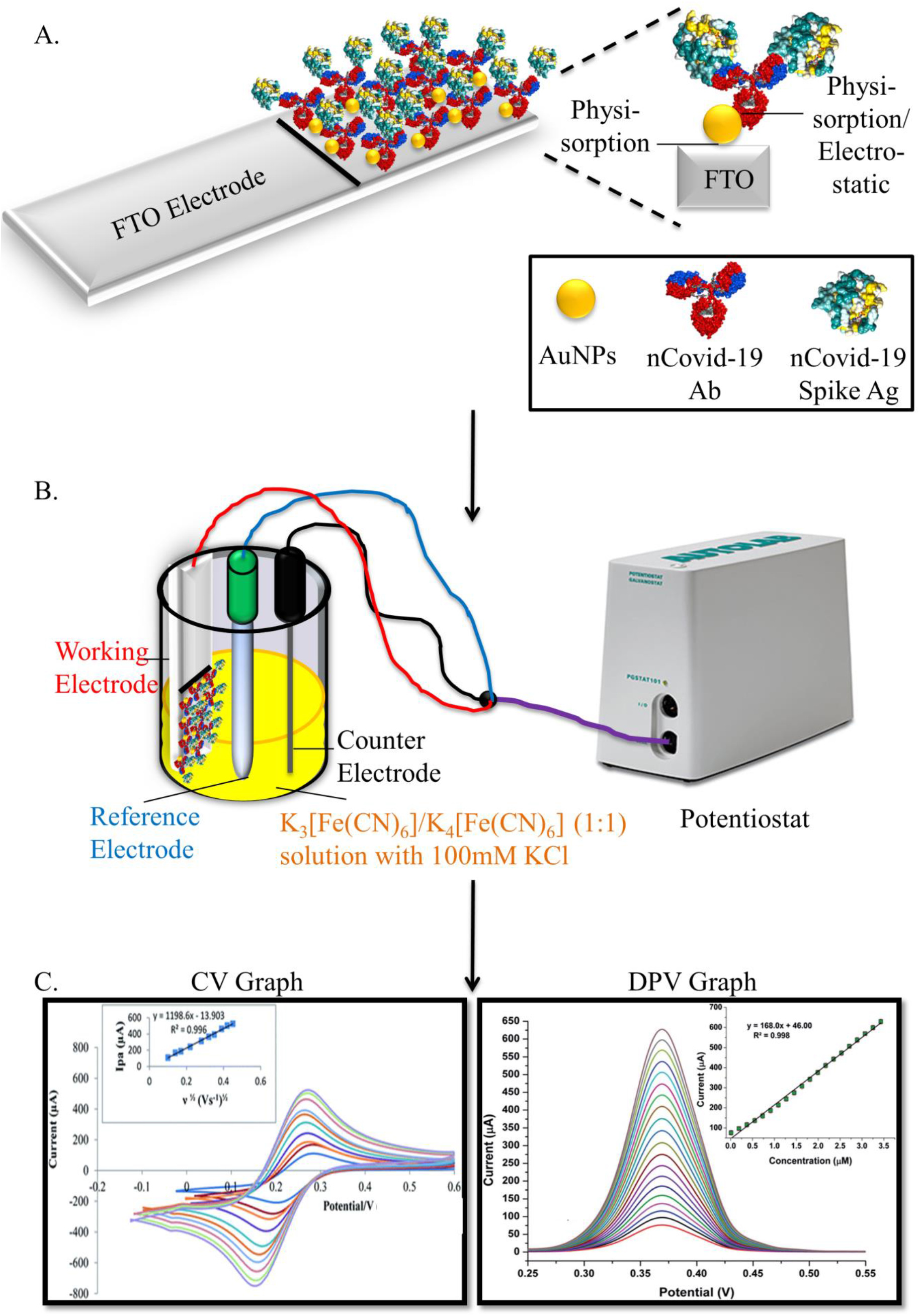
(A) The FTO electrode made up of a glass surface coated with fluorine doped tin oxide, consists of a sensing area which was first dipped in AuNPs colloidal solution to form an immobilised homogenous layer of AuNPs. Further, AuNPs are allowed to conjugate with nCovid-19 Ab either by physisorption or electrostatic bonding; (B) The fabricated electrode served as the working electrode in a 3 electrode system which consisted of a Ag/AgCl with KCl reference electrode and platinum counter electrode, immersed in K_3_[Fe(CN)_6_]/K_4_[Fe(CN)_6_] (1:1) solution with 100 mM KCl that acts as the redox potential buffer. The CV and DPV measurements were taken using a potentiostat. (C) The CV/DPV graph of each step of fabrication for detection at different concentration of nCovid-19 Ag showed changes in current that can be analysed to determine the LOD of fabricated FTO electrode.

### 3.2. Characterisation of AuNPs/nCovid-19 Ab labelling

UV spectra (Fig.1.(A)) showed the characteristic peak of AuNPs at 520 nm due to its surface plasmon resonance (SPR) properties whereas a red shift of 9 nm was observed at 529 nm when AuNPs were labelled with nCovid-19 Ab that confirmed the immobilisation of Ab on the surface of AuNPs via electrostatic interactions or physisorption mechanism. Three additional peaks were observed in FT-IR spectra (Fig.1.(B)) after conjugation of Ab with AuNPs at 1290 cm^−1^ (C-O stretching) and 2564 cm^−1^(S-H bond) and a new peak at 2328 cm^−1^ (C-N bond) confirmed the binding of AuNPs with nCovid-19 Ab. The change in hydrodynamic diameter was also observed from 21 nm (bare AuNPs) to 30±5 nm (AuNPs-Ab) further reconfirmed the conjugation of AuNPs with Ab (Fig.1.(C)) and the single and sharp peak showed that the particles are monodispersed in the colloidal solution. This increase in hydrodynamic diameter occurred due to binding of Ab with AuNPs. In Fig.1.(D), zeta potential was shifted from −42 mV (bare AuNPs) to −39 mV (AuNPs-nCovid-19 Ab) due to the insulating effect of the Ab protein deposition layer around the AuNPs which confirmed the process of conjugation. The size and morphological analysis of the AuNPs and AuNPs/nCovid-19 Ab conjugate was done by TEM as shown in Fig.1.(E)(i) & (ii) respectively. AuNPs were observed to be monodispersed with an average size of 21±5 nm while in the case of AuNPs/Ab, a proteinaceous layer can be seen deposited around the AuNPs verified the immobilisation of Ab with AuNPs.

**Fig.1.**
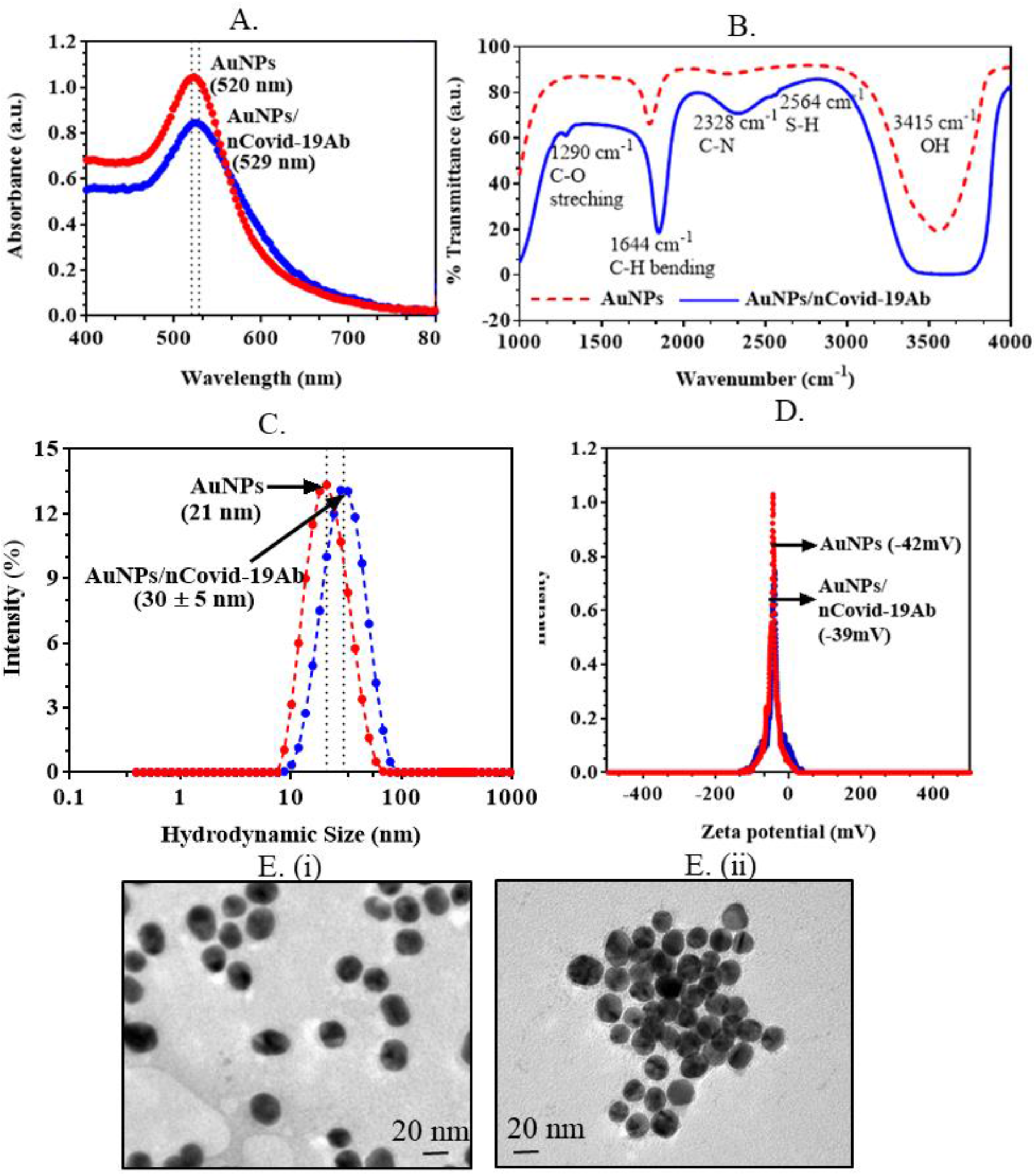
The labelling of nCovid-19 Ab with AuNPs: (A) The characteristic peak of bare AuNPs observed at 520 nm due to SPR whereas the peak broadened and showed a red shift to 529 nm when AuNPs labelled with Ab; (B) FTIR spectrum, three additional peaks observed after conjugation of Ab with AuNPs include two small peaks at 1290 cm^−1^ (C-O stretching) and 2564 cm^−1^(S-H bond) and a medium peak at 2328 cm^−1^ (C-N bond); (C) Hydrodynamic diameter increased from 21 nm in bare AuNPs to 30±5 nm in AuNPs/ Ab conjugate. (D) Zeta potential shifted from −42 mV (bare AuNPs) to −39 mV (AuNPs/ Ab) (E) TEM (i) monodispersed AuNPs with an average size of 20±5 nm and (ii) a proteinaceous layer of Ab observed deposited around the AuNPs.

### 3.3. Optimisation of fabricated FTO/AuNPs/nCovid-19 Ab

CV was carried out for optimisation of electrochemical parameters of FTO, FTO/AuNPs, FTO/AuNPs/nCovid-19 Ab and FTO/AuNPs/nCovid-19 Ab/Ag modified electrode as shown in Fig.2.(A). An increase in the current was observed between the bare FTO electrode and the FTO/AuNPs modified electrode possibly due to high conductivity of the AuNPs which accelerates the electron transfer on the surface of electrode (Tey et al., 2010). Additionally, high surface area of AuNPs allowed the binding of nCovid-19 Ab that was clearly indicated by increase in the redox peak for ferro/ferricyanide probe. This phenomenon could be explained due to increased permeation of ferro/ferricyanide probe that interacts with AuNPs and enhanced the conductivity. The increased current could be seen further when nCovid-19 Ag was applied on FTO/AuNPs/nCovid-19 Ab modified electrode. The analytical performance of the modified FTO/AuNPs electrode was optimized for nCovid-19 Ab where different concentrations of Ab (0.25 μg/mL, 0.5 μg/mL, 1.0 μg/mL, 1.5 μg/mL) were analysed and highest current output was observed in 1.0 μg/mL Ab concentration (Fig.2.(B)), that was further used in case of all electrochemical studies. The response time was evaluated from 5 s, 10 s, 15 s, 20 s, 25 s, 30 s and maximum peak current was observed at 25 s due to saturation of binding sites in nCovid-19 Ab with its spike Ag (Fig.2.(C)). CV data of the current output of the electrode at three different temperatures (4 °C, RT, 37 °C) (Fig.2.(D)) showed no effect of temperature on the functioning of the fabricated electrode hence all experiments were carried out at RT. Also, the optimum pH required to obtain maximum current signal of FTO electrode was determined by comparing CV at different pH range (4, 7.4, 8 and 12) and the maximum signal was observed at pH 7.4 as well as 8.0 (Fig.2.(E)). Therefore, buffer with pH 7.4 was used as the optimum pH for further detection experimentation.

**Fig.2.**
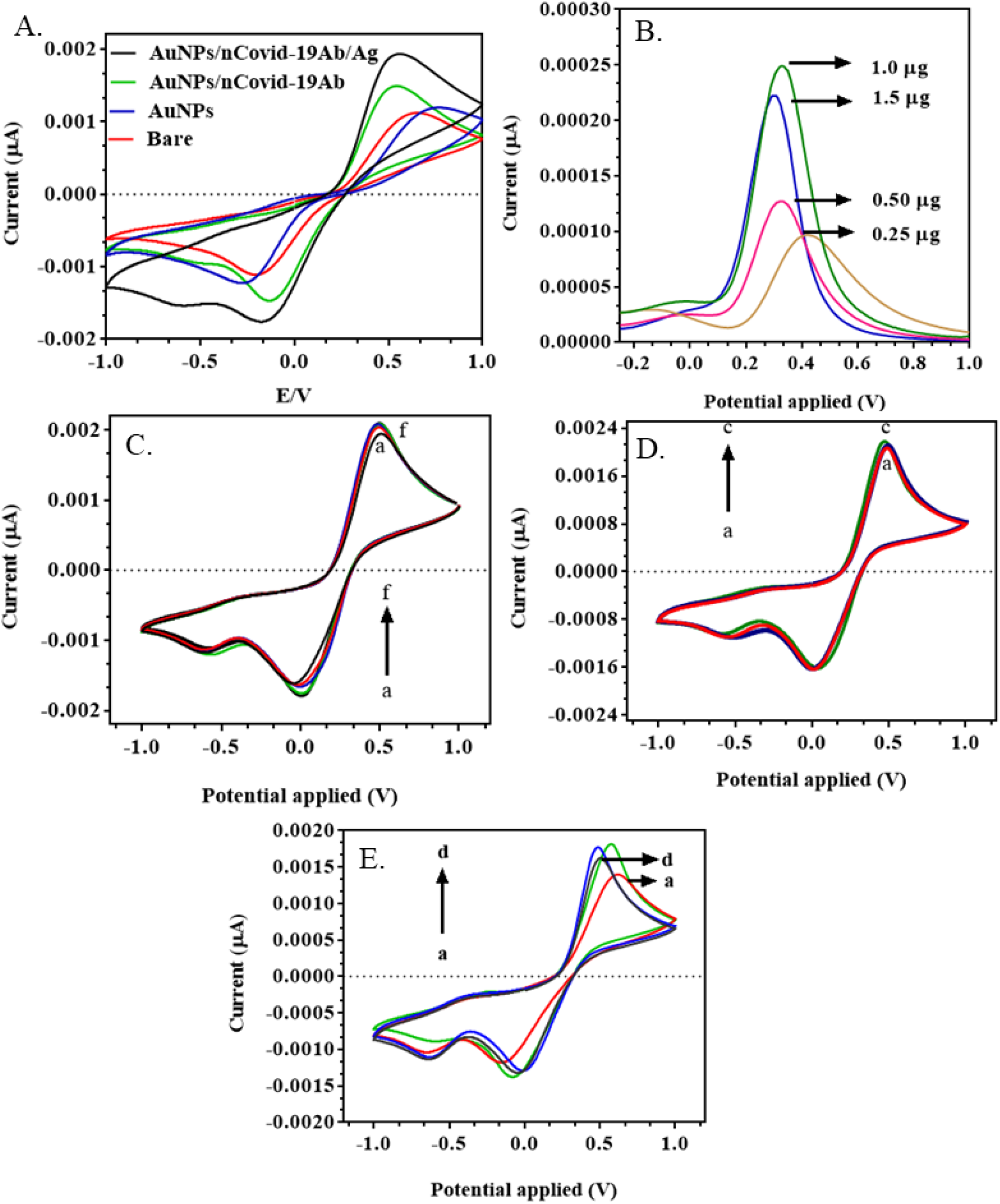
Electrochemical characterisation and optimisation of the fabricated FTO electrode in the scanning potential range of −1.0 V to 1.0 V in 100 mM KCl, having 0.005 M K_4_Fe(CN)_6_.H2O and 0.005 mM K_3_Fe(CN)_6_, at a scan rate of 100 mV/s: (A) CV spectra of bare FTO, FTO/AuNPs, FTO/AuNPS/nCovid-19 Ab and FTO/AuNPs/nCovid-19 Ab/Ag compared and it was observed that with the addition of each new biomolecule, the current output increased; (B) DPV of different concentrations of nCovid-19 Ab (0.25 μg/mL, 0.5 μg/mL, 1.0 μg/mL, 1.5 μg/mL) were taken and highest current output was observed in 1.0 μg/mL nCovid-19 Ab concentration; (C) CV spectra of different response time (a to f→5 s to 30 s) were superimposed and at 20 s and beyond the current output remained stabl; (D) CV at three different temperatures (a. 4 °C, b. RT, c. 37 °C) showed negligible change in current; (E) CV at different pH (a. 4, b. 7.4, c. 8, d. 12) where pH 7.4 and 8.0 inferred the highest current.

### 3.4. Analytical performance of the fabricated FTO/AuNPs/nCovid-19 Ab

Differential pulse voltammetry was used for the determination of nCovid-19 spike Ag concentration as shown in Fig.3.(A). The linear regression equation for DPV was explained in (Fig.3.(A)). For differential pulse voltammetry (DPV), the Intercept and slope was 0.002039 ± 6.396 e^−005^ and 0.0001784 ± 1.066 e^−005^ with equation I = 0.0001784× + 0.002039 (I (μA)) and r2 = 0.9722; where, I = peak current; c = concentration of nCovid-19 Ag. Different concentrations of nCovid-19 Ag ranging from 1 fM to 1 μM (standard buffer) were tested on the FTO/AuNPs/nCovid-19 Ab modified electrode and the standard calibration curve was plotted based on DPV (Fig.3.(B)) with LOD as 10 fM for nCovid-19 Ag. Cross reactivity studies were done to test non-specific binding of other viral Ag as shown in Fig.3.(C) that showed increased signal current with nCovid-19 Ag whereas no signal was observed with HIV, JEV or AIV Ag when all Ag concentrations were kept constant at 10 fM. This showed the high specificity of the nCovid-19 Ab towards nCovid-19 Ag. Furthermore, repeatability and stability parameters were also evaluated for FTO/AuNPs/nCovid-19 Ab fabricated electrode. In this case, readings of saliva samples spiked with nCovid-19 Ag were taken on a single modified electrode (Fig.3.(D)) with 120 fM concentration of nCovid-19 Ag. The results indicated that the fabricated electrodes can be used upto 3 times without major changes in peak current and could detect upto 120 fM. The stability of the FTO/AuNPs/nCovid-19 Ab fabricated electrode was observed at 7^th^, 14^th^, 21^st^ and 28^th^ day of its fabrication and kept at 4 °C. In Fig.3.(E) the modified FTO/AuNPs/nCovid-19 Ab electrode provide stable readings over a period of three weeks i.e. 21 days when tested over a period one month that shows the electrode can be stored at 4 °C upto 21 days and can be used for testing of nCovid-19 samples in laboratory set up for upto 3 weeks without any compromise in results.

**Fig.3.**
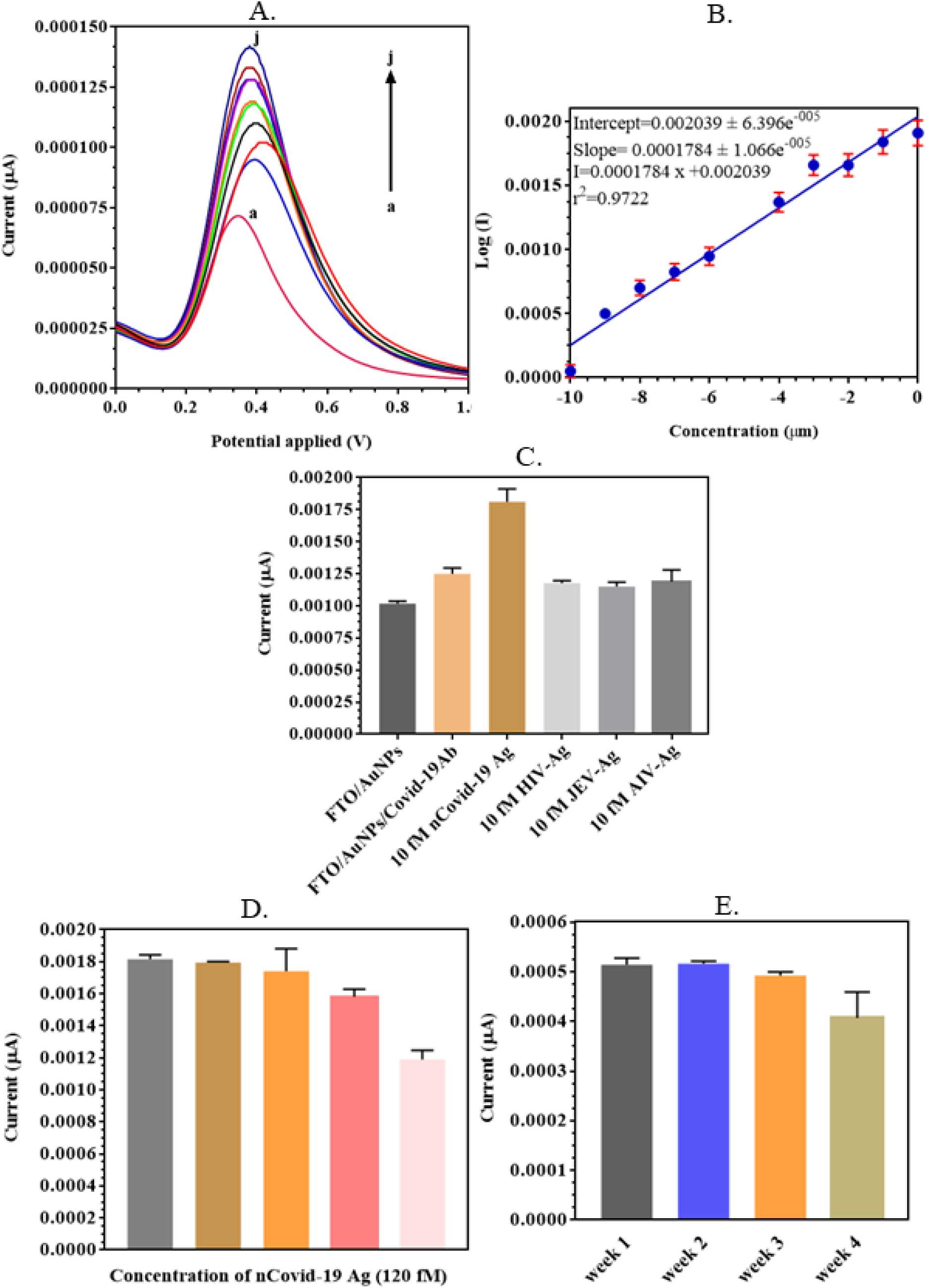
DPV of different concentrations of nCovid-19 Ag on the fabricated FTO/AuNPs/ nCovid-19 Ab electrode in the scanning potential range of −1.0 V to 1.0 V in 100 mM KCl, having 0.005 M K_4_Fe(CN)_6_.H2O and 0.005 mM K_3_Fe(CN)_6_, at a scan rate of 100 mV/s: (A) DPV of different nCovid-19 Ag concentrations (a) 1 μM (b) 100 nM (c) 10 nM (d) 1 nM (e) 100 pM (f) 10 pM (g) 1 pM (h) 100 fM (i) 10 fM (j) 1 fM in scanning potential range 0 V–1.5 V; (B) Standard calibration curve plot of the previous graph between log of the various concentrations of nCovid-19 Ag (a→j) and peak current; (C) Cross reactivity studies using HIV, JEV, and AIV Ag at fixed 10 fM concentration to check for specificity of FTO electrode; (D) Repeatability of the fabricated FTO/AuNPs/nCovid-19 Ab electrode tested on multiple saliva samples spiked with 120fM concentration of nCovid-19 Ag; (E) Stability of fabricated electrode tested upto the period of 1 month at 7 day time intervals.

### 3.5. Proof of principle of developed in-house novel electrochemical device eCovSens

The in-house built eCovSens device detects changes in the voltage by making use of an SPCE electrode that operates on the principle of signal transduction (Fig.4.(A)). The components of the modified biosensor include a bio-recognition element (nCovid-19 Ab), a transducer (carbon) and an electronic system composed of a display, processor and amplifier (in-house electrochemical instrument). The bio-recognition element, essentially a bio receptor, is allowed to interact with a specific analyte (nCovid-19 Ag) for detection. The Ag-Ab interactions interact with the transducer and provide a voltage signal as an output (Fig.4.(B)). The intensity of the signal output is proportional to the concentration of the analyte (nCovid-19 Ag). The signal is then amplified and processed by the electronic system and the output is converted from analogue to digital readings which are displayed on the device screen or mobile/computer if connected via bluetooth through an application which is possible due to the fabricated circuit (Fig.4.(C)).

**Fig.4.**
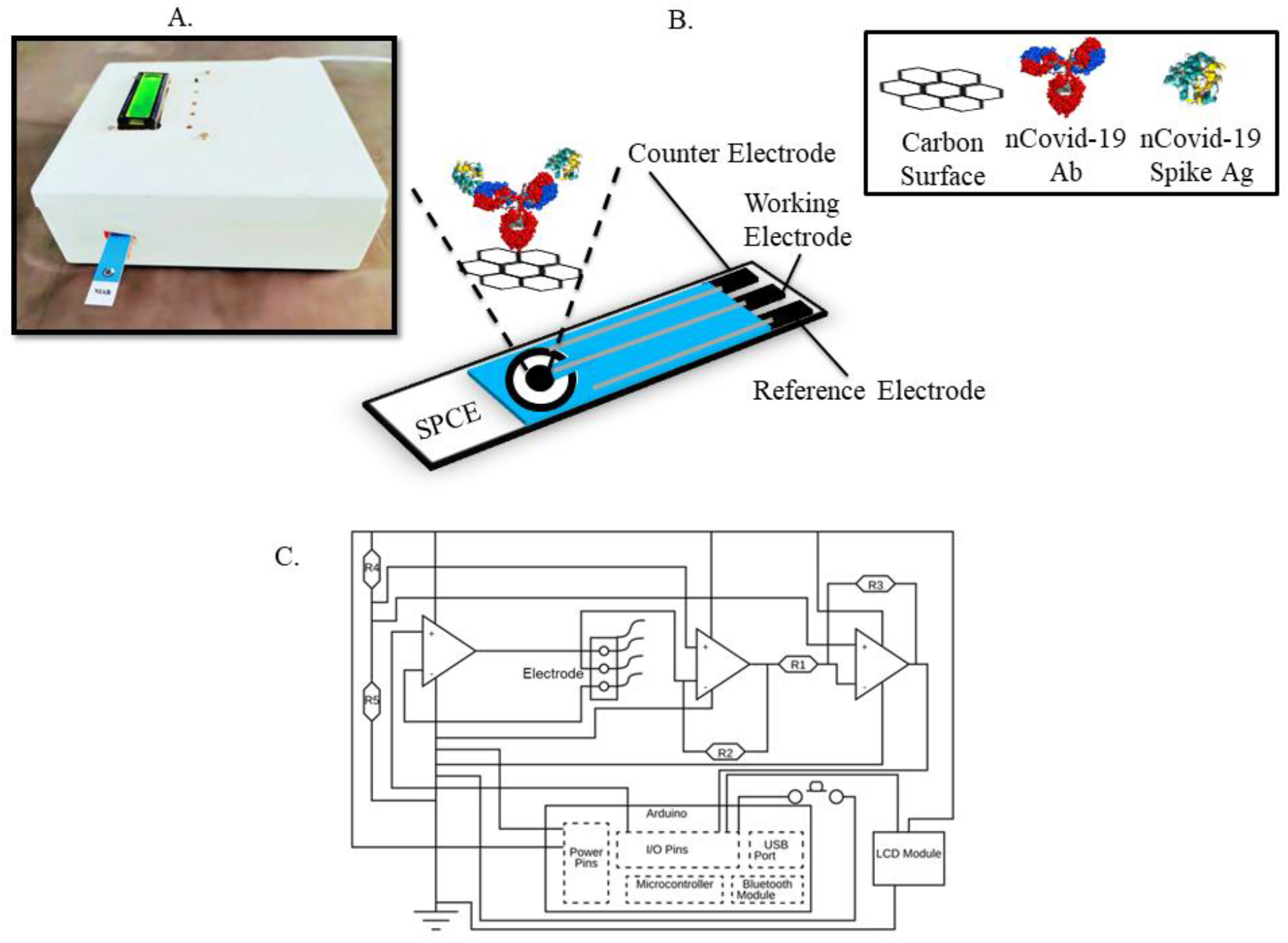
(A) The fabricated in-house built electrochemical eCovSens device; (B) Schematic of fabrication process of SPCE electrode where nCovid-19 Ab is allowed to immobilise onto the transducer of the SPCE followed by addition of nCovid-19 Ag and the transducer detects changes in electrical signal due to Ag-Ab interaction. (C) Circuit diagram of in-house built electrochemical device depicting various components and the connections.

### 3.6. Analytical performance of eCovSens

Analytical performance of the device was observed by immobilising nCovid-19 Ab at 1 μg/mL on SPCE electrode. Different concentrations of nCovid-19 Ag were prepared from 1 fM to 1 μΜ in 1× PBS, pH 7.4 and applied directly on the electrode. Change in voltage (in mV) was observed, wherein maximum signal was obtained at 100 nM concentration of nCovid-19 Ag and further increase in concentration (1 μM) did not alter the voltage signal. The device could read the output voltage within a rapid time of 1 min and detected upto 10 fM sensitivity (Fig.5.(A)), which is in close agreement with the commercially available potentiostat (Fig.3.(A)). No cross reactivity studies was observed with different concentrations of AIV Ag (Fig.5.(B)) which proved that the developed device is highly specific towards nCovid-19 Ag detection. Fig.5.(C) showed the change in voltage from 540 mV (control-normal saliva) to 638 mV (test-spiked nCovid-19 Ag (LOD-90 fM) in saliva) on fabricated SPCE/nCovid-19 Ab electrode. Fig.5.(D) shows the SPCE/nCovid-19 Ab electrodes were found to be stable with no change in voltage upto 4 weeks when tested with nCovid-19 Ag. Table 1 shows the currently in-use diagnostic techniques available for nCovid-19 detection which include molecular assays, immunoassays, and CT.

**Table 1.** Currently available diagnostic techniques for detection of nCovid-19.

| Type of Test | Institute | Limit of Detection | Reference |
| --- | --- | --- | --- |
| Virus blood culture and high-throughput sequencing of the whole genome | Wuhan Institute of Virology | Not Available | (Zhou et al., 2020) |
| Real time RT-PCR | Charité – Universitätsmedizin Berlin Institute of Virology | 3.9 copies per reaction for the E<sup>1</sup> gene assay<br>3.6 copies per reaction for the RdRp<sup>2</sup> assay | (Corman et al., 2020) |
| High Resolution CT (HRCT) | Huazhong University of Science and Technology | Not Available | (Pan et al., 2020b) |
| All-in-One Dual (DNA and RNA) CRISPR-Cas12a (AIOD-CRISPR) Assay | University of Connecticut Health Center | 1.2 copies DNA targets and 4.6 copies RNA targets | (Ding et al., 2020) |
| Rapid IgM-IgG combined Ab test kit | Guangzhou Medical University | Not Available | (Li et al., 2020b) |
| Closed tube one stage LAMP<sup>3</sup> | University of Pennsylvania | Not Available | (El-Tholoth et al., 2020) |
| Closed tube two stage isothermal amplification RAMP<sup>4</sup> assay | University of Pennsylvania | Not Available | (El-Tholoth et al., 2020) |
| RNA based paper LFA<sup>5</sup> PoC diagnostic device using LAMP assay | National Tsing Hua University | Not Available | (Yang et al., 2020) |
| ELISA<sup>6</sup> and gold immunochromatographic assay (GICA) for combined IgG-IgM | Wuhan University | Not Available | (Xiang et al., 2020) |
| Field Effect Transistor-based electrochemical biosensor | Korea Basic Science Institute | $1.6 \times 10^1$ pfu/mL | (Seo et al., 2020) |
| Potentiostat based immunosensor | National Institute of Animal Biotechnology | 10 fM | Current research work |
| eCovSens immunosensor | National Institute of Animal Biotechnology | 10 fM | Current research work |

<sup>1</sup>Envelope; <sup>2</sup>RNA-dependant RNA polymerase; <sup>3</sup>Loop-mediated isothermal amplification;
<sup>4</sup>Rapid analyte measurement platform; <sup>5</sup>Lateral flow assay; <sup>6</sup>Enzyme-linked immunosorbent assay

**Fig.5.**
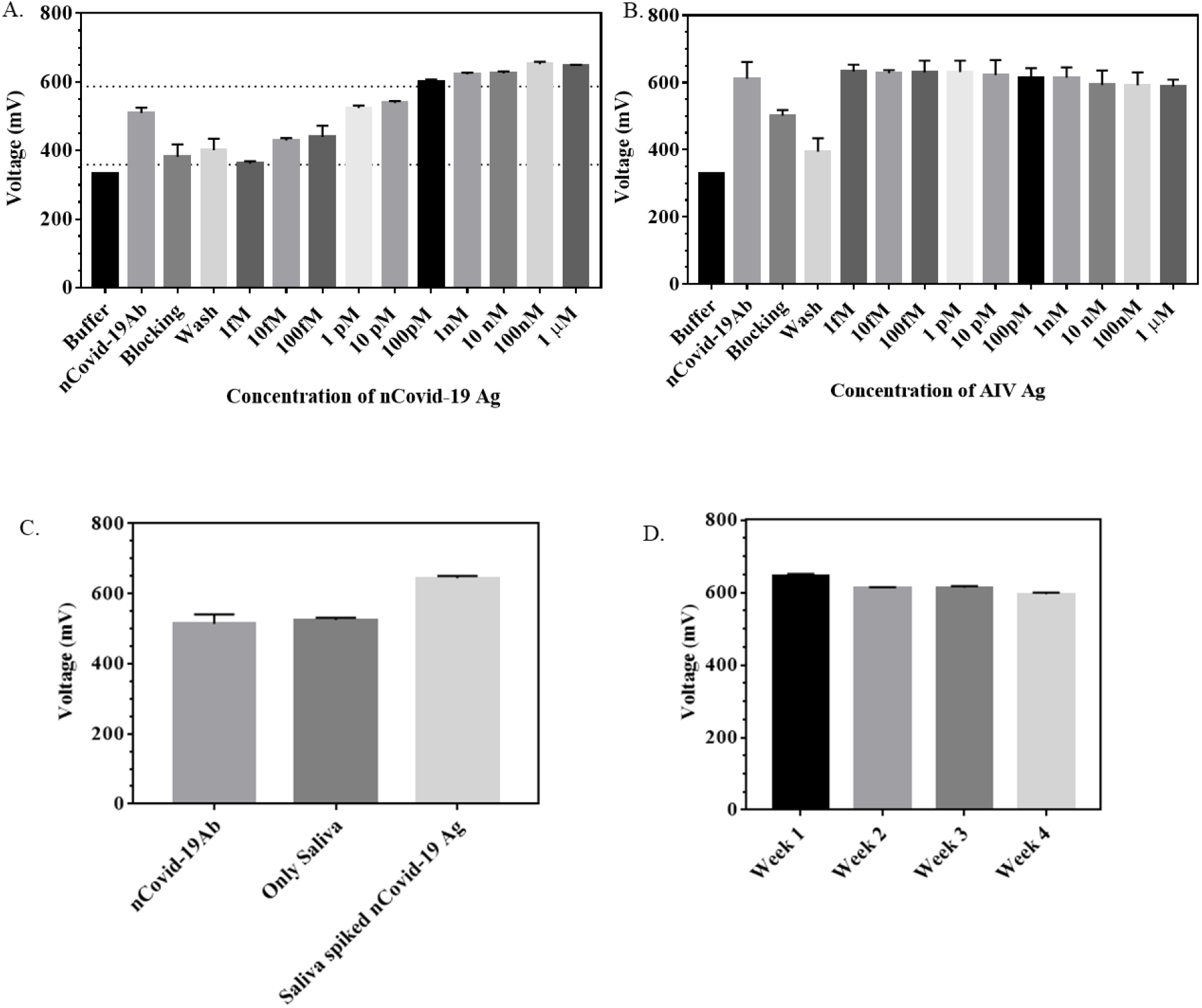
(A) and (B) represent output voltage data for different concentrations of Ag where (A) voltage due to specific binding of nCovid-19 Ab with respective nCovid-19 Ag while (B) represents nCovid-19 Ab binding with AIV Ag which shows no increase in voltage due to non-specific binding i.e. no cross reactivity; (C) Real sample analysis where saliva samples spiked with 90 fM of nCovid-19 Ag; (D) Stability of SPCE/nCovid-19 Ab electrode over a period of 4 weeks.

## 4. Conclusions

In the proposed work, we have successfully developed a novel in-house built electrochemical device eCovSens that can be used to detect changes in electrochemical signals upon Ag-Ab interaction on SPCE/nCovid-19 Ab modified electrode. The developed technology can be used to detect the global pandemic causing novel coronavirus nCovid-19 with LOD upto 10 fM within 10-30 s. We have also compared our findings with a potentiostat using an FTO/AuNPs/nCovid-19 Ab modified electrode with LOD upto 10 fM and found that our eCovSens device is equally sensitive in both buffer samples as well as spiked saliva samples along with no cross reactivity with other viral Ag, rapid detection, portable, and storage stability upto 4 weeks with no change in voltage. The device can further be potentially used directly on clinical samples for nCovid-19 detection. Moreover, unlike a potentiostat, this device is portable since it can be battery operated as it uses very less voltage (1.3-3V), and can be used for bedside/on-site PoC diagnostics. The developed device is very cheap and cost effective as compared with the commercial potentiostat which is an expensive instrument that requires a laboratory set up. The in-house built novel electrochemical device shows great future potential for detection of various other diseases as the sensor can easily be customised.

## Acknowledgements

We acknowledge the Department of Biotechnology (DBT), New Delhi, for the grant BT/AAQ/01/NIAB-Flagship/2019 and C0038 as internal core research support from DBT-National Institute of Animal Biotechnology (DBT-NIAB), Hyderabad. A.R. would like to acknowledge DST-INSPIRE fellowship (IF180729) sponsored by Department of Science and Technology (DST), New Delhi.

